# Temporal changes in genetic variation during the invasion of *Lygodium japonicum* in North America

**DOI:** 10.64898/2026.11.05.764610

**Authors:** Jessie A. Pelosi, Abby R. Pearse, Poppy C. Northing, Morgan L. Markley, Ethan Altergott, James B. Beck, Katrina M. Dlugosch

## Abstract

**Premise of the study:** Invasive species are now widespread and have detrimental ecological and economic impacts. They also offer opportunities to investigate rapid evolution in colonizing populations. Here, we leverage nuclear DNA from herbarium specimens of *Lygodium japonicum*, an invasive fern to identify the putative source(s) of its invasion, 2) test for climate adaptation in its invaded range, and 3) characterize the effects of genetic bottlenecks on adaptive and neutral variation during its establishment.

**Methods:** We use low-coverage genome skimming data from specimens spanning 1905 through 2017 to identify the source region(s) of the *L. japonicum* invasion of North America. We use a genotype-environment analysis of invaded range samples to identify genomic regions associated with local adaptation to variation in five bioclimatic variables. Finally, we compare heterozygosity between native and invasive individuals and their changes through time in non-coding, coding, and putatively adaptive loci.

**Key results:** We identify mainland Asia as the most likely source region for this invasion. Our genotype-environment analysis found 144 candidate SNPs significantly associated with climatic variation in the invaded range, with precipitation of the driest month associated with the greatest number of SNPs, suggesting rapid genomic adaptation in the invaded range. Finally, we found conflicting trends in individual-level heterozygosity over time when analyzing different types of genetic variation: in non-coding loci, there was a small, yet significant decrease in genetic diversity, but in putatively adaptive loci we did not find evidence of a genetic bottleneck.

**Conclusions:** Natural history collections provide exceptional opportunities to explore evolutionary processes, by capturing genomic sequences across time. Here, herbarium specimens reveal rapid evolution, maintenance of adaptative variation, and accumulation of noncoding diversity in the *Lygodium japonicum* invasion, highlighting the potential for adaptative variation to fuel the success of invasive species.

## INTRODUCTION

Anthropogenic activity has led to dispersal of species into novel environments; in some cases, these introduced species become invasive by forming self-sustaining, ecologically-dominant populations that outcompete native flora (Mack et al., 2000; van Kleunen et al., 2015). The negative ecological impacts of species invasions are wide-ranging and include community homogenization (Olden et al., 2004; Vilà et al., 2011; Petsch et al., 2022), altered nutrient cycling (Vilà et al., 2011; Pyšek et al., 2020), disrupted disturbance regimes (Ehrenfeld, 2010; Pyšek et al., 2020), and introductions of novel pathogens via hitchhiking (Brasier, 2008; Burgess et al., 2016; Burgess and Wingfield, 2017). In addition, invasive species pose unprecedented economic impacts to both natural and agricultural lands (Diagne et al., 2021; Fantle-Lepczyk et al., 2022). It is estimated that, by 2017, costs related to invasive species had reached over $1 trillion globally (Diagne et al., 2021). Understanding how invasive species successfully colonize novel environments is essential to mitigating these impacts and managing their spread in natural and cultivated ecosystems (Allendorf and Lundquist, 2003; Matheson and McGaughran, 2022).

At the same time, species invasions offer exceptional opportunities to understand fundamental evolutionary processes that occur on ecological timescales (Baker and Stebbins, 1965; Lee, 2002; Prentis et al., 2008). The initial introduction, subsequent establishment, and spread of an invasive species represent three distinct periods along the “invasion continuum” (Richardson et al., 2000). Identifying the introduction source(s) of invasions is critical for management purposes (van Kleunen et al., 2015; Buckley and Catford, 2016), but also for making appropriate genomic, trait, and ecological comparisons between native and invading populations (Dlugosch and Parker, 2008). This enables investigations that can accurately determine whether traits and genetic variants differ in invasions, as well as whether observed phenotypic differences between native and invading populations arise from standing genetic variation in the native range or arose *de novo* (Hodgins et al., 2018, 2025). A combination of historical records and genetic data can be used to elucidate the initial source(s) of an introduction (e.g., Hendry, 1931; Barker et al., 2017).

Subsequent to their introduction, invading populations may need to adapt to novel environmental conditions in order to establish self-sustaining populations which can spread and become ecologically dominant (Richardson et al., 2000; Colautti and MacIsaac, 2004; Blackburn et al., 2011). Despite demographic bottlenecks that commonly occur during an introduction of an invader (Sakai et al., 2001), previous work on the population genomics of invaders has found little evidence that genetic bottlenecks are typically severe or limit the adaptive potential of invaders (Dlugosch et al., 2015). Indeed, evidence suggests that invasive populations are capable of rapid adaptation to their invaded range (North et al., 2021; Hodgins et al., 2025), and can rapidly adapt to local environmental conditions (Oduor et al., 2016). Outstanding questions remain about the nature of how introduction history shapes an invasion (Colautti et al., 2006; Bucharova and van Kleunen, 2009), the kinds of genomic variation that facilitate invasions and drive invasion success (Dlugosch et al., 2015), and the degree of parallelism in the genetic mechanisms involved in adapting to invaded environments (Battlay et al., 2025; Hodgins et al., 2025).

Natural history collections (e.g., herbarium specimens) are valuable resources that serve as time capsules that enable research into processes that occur on relatively short timescales, such as species invasions (Lang et al., 2019; Sutherland et al., 2021; Burbano and Gutaker, 2023; Kim et al., 2023). Museum specimens harbor a myriad of information that is pertinent for investigating ecological and evolutionary phenomena, including biogeographical data from metadata (e.g. Chauvel et al., 2006; Lankau et al., 2009), phenotypic data (e.g., morphological, physiological, chemical, or phenological traits; Heberling, 2022), species interactions (Meineke et al., 2018a; Ristaino, 2020), and genomic DNA preserved in specimen tissues (Meineke et al., 2018b; Latorre et al., 2020; Burbano and Gutaker, 2023). Sequencing genomic DNA of individuals collected throughout the duration of invasions enables temporal investigations into population genomics of the introduction, spread, and establishment of invaders (Kim et al., 2023; McGaughran et al., 2024). Previous studies have used genomic chronosequences from historical specimens to elucidate invasion history (Martin et al., 2013, 2014; Yoshida et al., 2013; Bradshaw et al., 2021; Bieker et al., 2022), how hybridization influences colonization (Bock et al., 2021; Bieker et al., 2022), the role of new mutations in colonization (Exposito-Alonso et al., 2018), and the genetic basis of traits putatively involved in rapid adaptation to novel conditions in the introduced environment (Vandepitte et al., 2014; Alves et al., 2019; Kreiner et al., 2022; Stuart et al., 2022).

Ferns, the second largest group of vascular plants, make up non-trivial proportions of non-native flora (Pelosi, Yañez et al., in press; Jones et al., 2019), but they are largely understudied. One such invasive fern is *Lygodium japonicum* (Thunb.) Sw., a prolific vine introduced to the southeastern United States in the late nineteenth and early twentieth century through the horticultural trade (European and Mediterranean Plant Protection Organization, 2019). *Lygodium japonicum* is a member of one of only two clades of ferns that have a vining habit, which is facilitated by indeterminate rachis growth and dormant pinnae buds (Mueller, 1982, 1983). Like many *Lygodium* species, *L. japonicum* can form dense ground cover, sprawling mats, and climbing fronds that can negatively impact native ecosystems by outcompeting native taxa (Leichty et al., 2011) and altering natural fire regimes (Ferriter, 2001; Koop, 2009). Interestingly, extensive proliferation of these mats in the invaded range is less common, which is possibly related to seasonal freezing (Zeller and Leslie, 2004). This species mostly inhabits mesic and hydric environments in its native range, although it can occupy a wide variety of conditions including xeric pine forests in the invaded range (Van Loan, 2006). It has quickly spread into natural and agricultural lands (e.g., pine plantations) throughout most of the southeastern United States (Graves, 1920; Matthews, 1940; Diddell, 1941) (Fig. 1). Despite being part of the horticultural trade, nearly nothing is known about the origin source(s) of these plants. Although Hutchinson and Langeland (2010) suggest that multiple source populations are likely, they do not provide a rationale for this inference. Once established, further spread was likely facilitated by the movement of minute airborne spores (thousands of which can be produced by a single leaflet), and contaminated pine-straw mulch (Zeller and Leslie, 2004).

**Figure 1.**
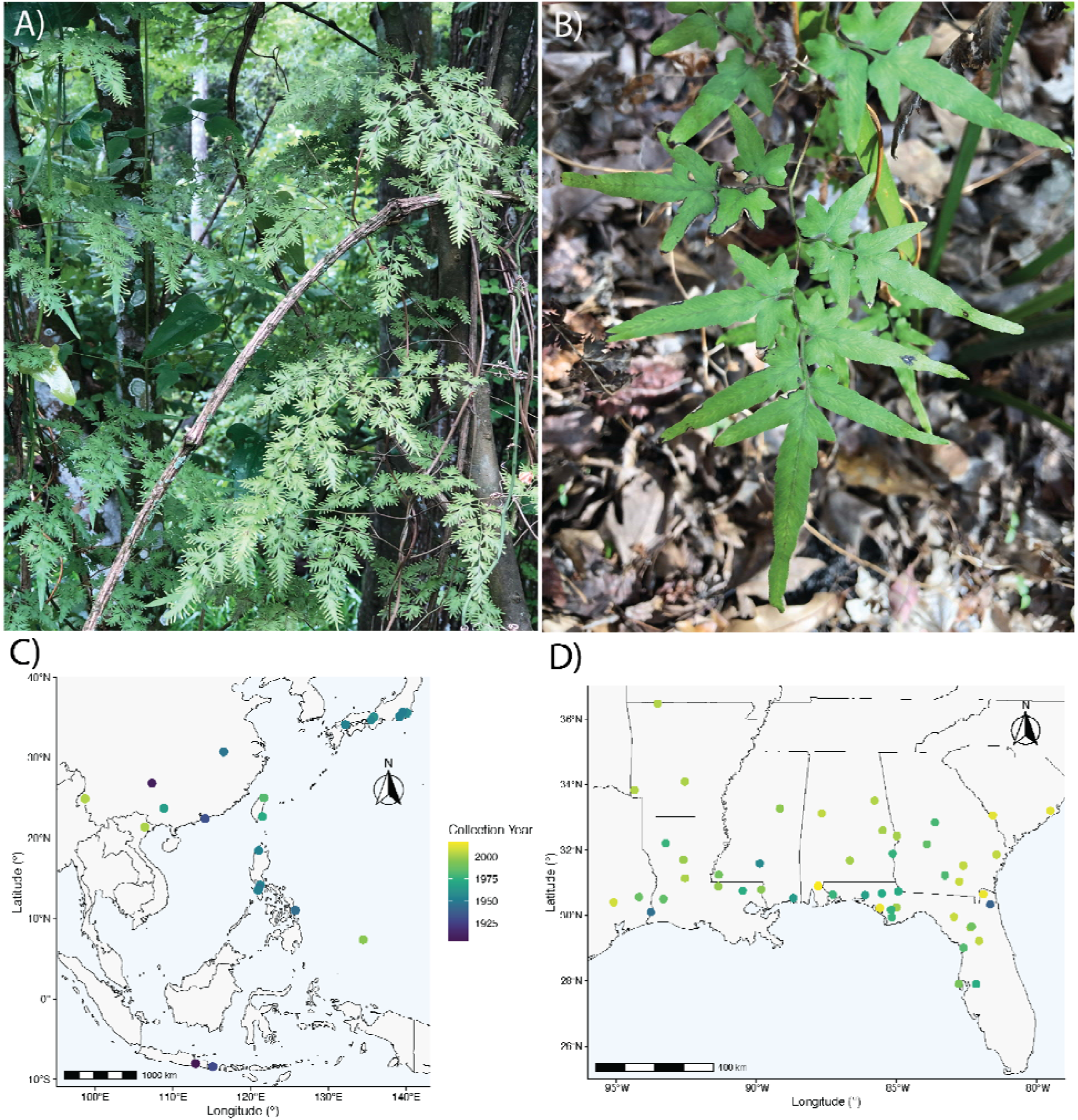
Photographs of *Lygodium japonicum* from the invaded range (J.A. Pelosi). (A, B). Sampling locations of specimens in the native (right, C) and invaded ranges (left, D). Symbols are colored according to collection year.

The *L. japonicum* invasion offers a valuable opportunity to explore the evolutionary processes facilitating the colonization of non-native taxa across ecological timescales. In particular, genomics of herbarium specimens could provide insight into the origin(s) of the invasion, whether populations in the invaded range are evolving, and whether natural selection imposed by novel environmental pressures are driving rapid adaptive evolution in this invasion. Markley et al. (2024) generated genome-skimming data from herbarium samples spanning 112 years from the native and invaded range, the first genomic dataset to investigate evolutionary processes in the *L. japonicum* invasion. Their analysis focused on the chloroplast genomes sequenced in these data, from which they found that the native range samples (N = 24) contained 17 haplotypes while the invaded range samples (N = 50) contained only three: one dominant haplotype across 47 invader samples and two additional haplotypes among three invaders. This suggested a potential loss of genetic diversity in the invasion, but also putative evidence of multiple introductions (contributing different chloroplast lineages), though both of these inferences were tentative given that plastid variation does not reflect the bulk of genomic variation within a species (Smith, 2015). Markley et al. (2024) also found that the dominant haplotype from the invasion was not among the samples from the native range, but differed by a single mutation from a haplotype observed in China. The only haplotype shared between ranges was a rare haplotype from the invasion, which was also observed in China and the Philippines. These results are consistent with prior introduction(s) from China, but are not definitive without observations of the relationship of invading genotypes to those in the native range.

In the present study, we expand on the work of Markley et al. (2024) by analyzing nuclear genomic variation in their samples spanning 112 years. We leverage their sequence data to identify polymorphisms across the genome and use an available *L. japonicum* transcriptome (Qi et al. 2018; Pelosi et al. 2022) to categorize these polymorphisms as occurring in coding vs. non-coding genomic regions. We further investigate whether these polymorphisms show evidence of being involved in local adaptation using genome-environment association analyses. Using the greater resolution of these nuclear data, we ask: 1) Which is/are the source region/s of the *L. japonicum* invasion in the southeastern U.S.? 2) Is there adaptation to environmental variation during the expansion of *L. japonicum* across its invaded range? And 3) How does any genetic bottleneck experienced during establishment by the invading population of *L. japonicum* affect putatively adaptive variation and the potential for adaptation to have aided its establishment?

## METHODS

### Data acquisition and processing

Raw whole-genome skimming sequence data generated by Markley et al. (2024) were accessed from NCBI BioProject PRJNA1114707 (mean number of bases 3.07Gb and coverage ∼0.21x per sample based on an estimated haploid genome size of 14.53 Gbp (Pelosi et al., 2023). Sequence data was generated from DNA extracted from 74 herbarium specimens of *L. japonicum* that spanned its native range (N = 24; 1905-2003; Japan, China, Vietnam, Indonesia, Taiwan, Palau, Philippines) and introduced range (N = 50; 1940-2017; United States: Florida, Georgia, South Carolina, Alabama, Louisiana, Arkansas, Mississippi, Texas) (Fig. 1; Table S1). For details on sampling and data generation, see Markley et al. (2024). TrimGalore v0.6.10 (https://zenodo.org/records/7598955) was used to remove low-quality reads and adaptors from the raw sequencing data. Reads with quality scores greater than Q20 and length greater than 30bp were retained (--quality 20 --length 30).

### Genomic reference

A reference genome for *L. japonicum* is not yet available. Therefore, we assembled a *de novo* reference sequence using the trimmed reads from Markley et al. (2024) using CAPTUS (Ortiz et al., 2026), a platform designed for the assembly of high-throughput sequencing data generated by targeted-capture, genome skimming, whole genome sequencing, and RNASeq. Trimmed reads were passed to the ‘assemble’ module of CAPTUS, which uses MEGAHIT (Li et al., 2015) to generate assemblies for each sample and salmon (Patro et al., 2017) quantify read coverage and filter out low-coverage contigs, with the ‘CAPSKIM’ preset parameters, Then we used the ‘extract’ module to cluster contigs at 79.2% sequence identity with MMSeqs2 (Steinegger and Söding, 2017) requiring at least 20% of samples present for a cluster to be retained. The resulting assembly consisted of 71.8 Mbp across 66,096 contigs with a contig N50 of 1.1 Kbp. We refer to this reference as ‘LJCAPTUS.’

To differentiate between coding and non-coding loci, we used BLASTx (Camacho et al., 2009) to determine sequence homology between the LJCAPTUS assembly and an available *L. japonicum* transcriptome (Qi et al., 2018; Pelosi et al., 2022; available: https://github.com/jessiepelosi/ferntxms/tree/main/Data/Assemblies, specimen code ‘LYJA’), using an E-value threshold of 1 × 10^−5^. We then subsetted LJCAPTUS contigs based on these BLAST results: loci with BLAST hits to the transcriptome were classified as coding, while all other loci (e.g., no significant BLAST hits to the transcriptome) were classified as non-coding. Unless otherwise noted, the entire set of loci (coding and non-coding) were combined for analyses.

### Alignment and genotyping

Trimmed reads were mapped to LJCATPUS with bwa (Li and Durbin, 2009), and flagstat in samtools v1.19.2 (Li et al., 2009) was used to assess mapping rates. Optical and PCR duplicates were marked with MarkDuplicates in GATK v4.2.5.0-9 (McKenna et al., 2010). Genotypes were called following GATK BestPractices (Van der Auwera et al., 2013). Briefly, read groups were added with picard v2.23.4 (Broad Institute, 2019), initial variant calling was done with HaplotypeCaller in GATK (McKenna et al., 2010), and initial variant calls were combined using GenomicsDBImport in GATK v4.2.5.0-9 (McKenna et al., 2010). Given that *L. japonicum* is tetraploid in at least part of its range (Pelosi et al., 2023, 2024 and references therein), we treated every individual as tetraploid with the flag ‘-ploidy 4’ in the HaplotypeCaller command. Final genotype calls were produced using GenotypeGVCFs (using flag ‘--sample-ploidy 4’) in GATK v4.2.5.0-9 (McKenna et al., 2010).

From these genotyped variants, only SNPs were retained using GATK SelectVariants. Genotypes with a depth < 10 and genotype quality < 20 were marked as missing data using GATK VariantFiltration. Loci with more than 50% missing data were removed with bcftools v1.15.1 (Danecek et al., 2021). Given that DNA from herbarium samples may undergo de-amination (Weiß et al., 2016; McAssey et al., 2023) such that cytosines are converted to uracil, we removed heterozygous C-to-T variants, which may correspond to degradation and not true genetic variation, with the +setGT plug-in in bcftools v1.15.1 (Danecek et al., 2021). Minor allele frequency (MAF) was then re-calculated for every site with the +fill-tags plug-in in bcftools v1.15.1 (Danecek et al., 2021). Sites with MAF > 0.05 were retained and only one variant was retained per contig to avoid biasing analyses with linked sites.

### Population structure

To investigate patterns of population structure in the invasion, we explored relationships among individuals with an ordination-based approach. Missing genotypes were imputed by replacing missing data with mean allele counts with the R package poppr v2.9.8 (Kamvar et al., 2014). The dudi.pca function in ade4 v.1.7.23 (Dray and Dufour, 2007) was used to conduct a principal component analysis (PCA) based on the genomic data, from which the first 50 principal components were retained. Significant associations between genetic distance and geographic distance of individuals (i.e., isolation by distance, IBD), and between genetic distance and temporal distance of individuals (i.e., isolation by time, IBT) were tested for within each range (native and invaded). Pairwise genetic distances were calculated using the Prevosti’s method (Prevosti et al., 1975) with poppr v2.9.8 (Kamvar et al., 2014). Euclidean geographic and temporal distances were calculated with stats v4.4.1 (R Core Team, 2025). Mantel tests with 999 permutations were used to determine if there were significant correlations between genetic, geographic, and temporal distances. All analyses were done in R v4.5.3 (R Core Team, 2025).

### Genetic distance of invaded range plastid haplotypes

Three plastid haplotypes were identified by Markley et al. (2024) in the invaded range of *L. japonicum*. Nearly all samples (47) in the invaded range had a single chloroplast haplotype; the other two invaded range haplotypes were represented by either one (haploptype #2 from 1965) or two samples (haplotype #3 samples from 1996 and 1999). To assess whether these haplotypes might also be reflected in the nuclear data, we compared the distribution of the median genetic distances between samples within the dominant plastid haplotype with the median genetic distance of the samples in the other plastid haplotypes with the samples in the dominant plastid haplotype using Prevosti’s method as above. We then used a percentile rank via empirical cumulative distribution function test to compare the genetic distance between the different haplotypes with the background distribution.

### Genomic loci associated with local adaptation to climate

A genotype-environment association (GEA) analysis was used to identify loci where there were significant correlations between allele frequencies and climatic variables. Only invaded range samples were included in this analysis. Geographic coordinates were estimated for each sample based on locality information on the associated herbarium labels (Table S1). Temperature and precipitation data were downloaded for each year from 1901-2025 from the Climatic Research Unit gridded Time Series (CRU TS; Harris et al., 2020). Nineteen bioclimatic variables were extracted for each sample in its corresponding collection year with dismo v1.3-16 (Hijmans et al., 2024). Each of the 19 bioclimatic variables were visually examined to determine if gradients across the invaded range were present. Only bioclimatic variables that were not highly correlated (i.e., R^2^ value < 0.8) were retained; for highly-correlated variables we retained the one with a greater loading in a PCA of the 19 total bioclim variables. Each of our selected bioclimatic variables was visualized using a PCA to observe the loading of each bioclim variable. We retained five bioclimatic variables for our GEA that were both independent and strongly loaded on the major PC axes: annual mean temperature (BIO1), mean diurnal range (BIO2), precipitation of the wettest month (BIO13), precipitation of the driest month (BIO14), and precipitation seasonality (BIO15). Each bioclimatic variable was mapped against the invaded range to visualize where samples are located against the environmental gradient (Supplemental Figure 1).

A partial redundancy analysis (pRDA) GEA, a multivariate approach used to detect significant associations of allele frequencies and environmental variables, was performed using the SNP dataset with the LJCAPTUS reference. The pRDA was conditioned on the first two PC axes of a genomic PCA to account for population structure. Candidate loci were categorized as those outside of three standard deviations from the mean loading value on the first three RDA axes. The number of candidate SNPs for each environmental predictor were totalled (Kamvar et al., 2017). Significant differences in the proportion of candidate SNPs explained by each environmental predictor in coding versus non-coding loci were evaluated using a Chi-squared test with “chisq.test” in stats v4.4.1 (R Core Team, 2025).

### Genetic variation over time

To assess if and how genetic variation in both the native and invaded range changed over time, observed individual heterozygosity was calculated for four sets of loci using dartR v.1.0.7 (Gruber et al., 2018): 1) the total set of loci, 2) coding loci, 3) non-coding loci, and 4) climate-associated loci identified from the pRDA. To account for tetraploidy, the function gl.report.polyploid_heterozygosity function in dartR v1.0.7 (Gruber et al., 2018) was used. Linear models were fit to these data to analyze whether there were significant changes in individual observed heterozygosity over time. First, models were fit with fixed effects of collection year (standardized to the first year in the dataset, 1905), invasion status, and an interaction term of collection year and invasion status as predictors of observed heterozygosity. Given that main effects can be difficult to interpret with significant interaction terms, linear models were fit for each range separately if the interaction term was significant. If the interaction term was not significant, a reduced model was fit without the interaction of collection year and invasion status.

## RESULTS

### Dataset Assembly

Mapping rates to the LJCAPTUS reference sequence were moderately high, with an average of 71.46% of reads mapped per sample (range: 40.45 - 80.41%). After filtering for low-quality and low-depth sites, a total of 4,291 variant loci were retained. Out of these variant loci, 1,624 were coding and 2,667 were non-coding.

### Population Structure

PCA of the genomic data showed clear clustering of the invaded range samples (Fig. 2), and a more concentrated cluster of native range samples closer to samples from earlier in the invasion (Fig. 2c, d). Indeed, there was a significant, but weak, linear relationship between the date of collection of invaded range samples and their median genetic distance to the native range samples (F_1,47_, P = 0.013, Adj. R^2^ = 0.1055). There was no clear clustering geographically across the invaded range, although there did appear to be clustering temporally with samples from later in the invasion (more recent) occupying a greater proportion of genomic space (Fig. 2) distinct from samples collected earlier and the native range. In the native range, samples from Palau and Taiwan were genomically distinct from other samples in the native range. After removing these samples, there was no distinct geographic clustering between native and invaded samples.

**Figure 2.**
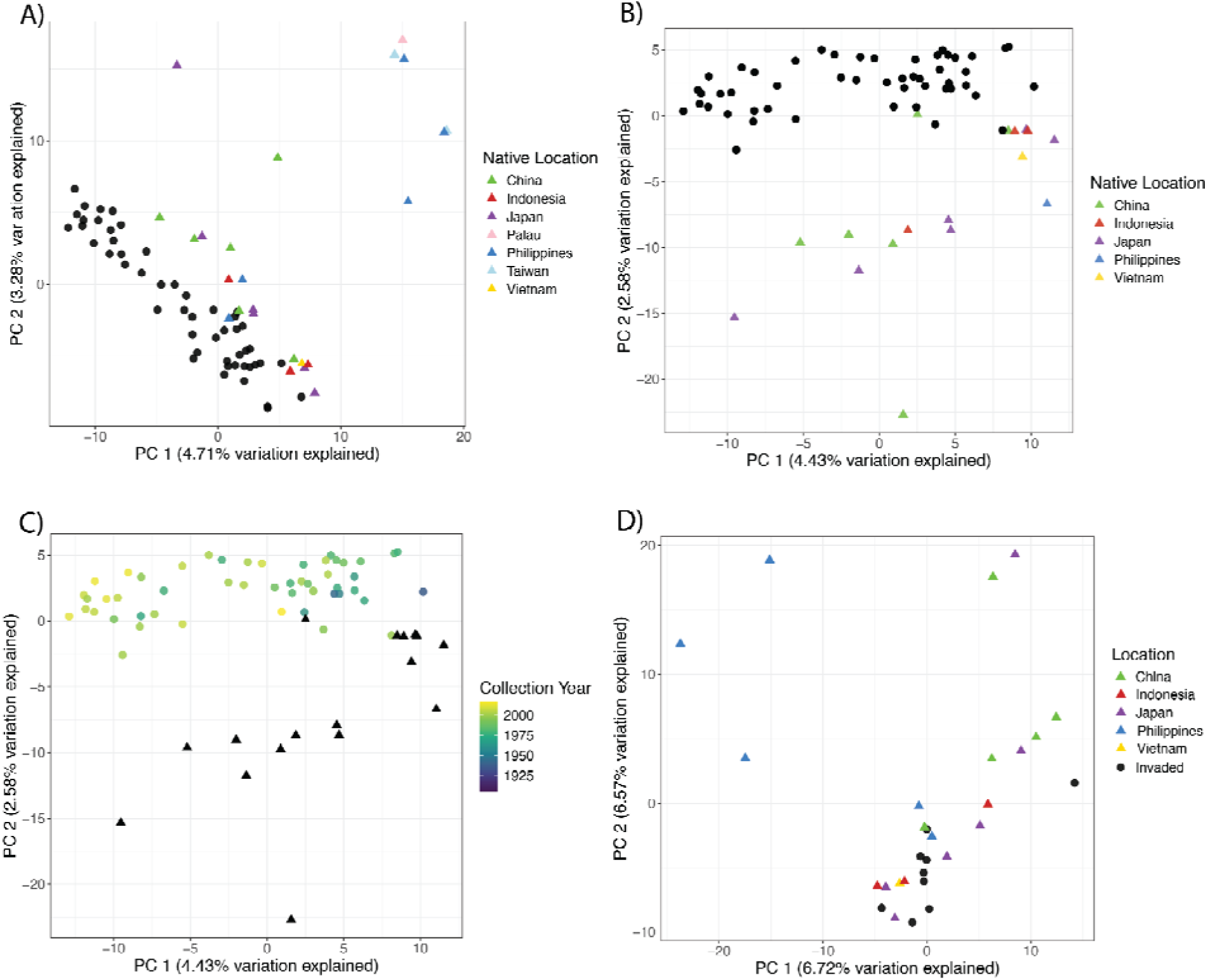
Genomic principal component analysis (PCA) of *Lygodium japonicum* samples used in this study. (A) and (B) depict geography of the native range (colored triangles) as it relates to the invaded range (black dots). In C, samples from the invaded range (dots) are colored based on their collection year. In D, samples are colored as in A and B, but only early invasion (pre-1975) samples are shown. B, C, and D are PCAs after removing three outlying samples from two island nations (Taiwan and Palau).

There was not a significant correlation between genetic and geographic distance in the invaded range (R = -0.1370, p = 0.947), nor in the native range (R = -0.1260, p = 0.778; Table S2). Furthermore, there was no significant correlation between genetic and temporal distance in the invasion (R = -0.2761, p = 0.995) nor the native range (R = -0.0282, p = 0.601; Table S3).

When comparing the genetic distance between samples within the invaded range among the three different chloroplast haplotypes identified by Markley et al. (2024), the genetic distances of samples in the alternative plastid haplotypes (#2 and #3) were not significant outliers compared to the distribution of the genetic distances within the dominant plastid haplotype (percentile rank via empirical cumulative distribution function, all *P* > 0.05) (Supplemental Figure 2).

### Identification of Climate-Associated Loci

Samples from the native range tended to occupy areas that are wetter (mean precipitation of the wettest month = 398.95 mm, range: 170.30 - 780.20 mm; mean precipitation of the driest month: 32.69 mm, range: 0.20 = 177.30 mm) compared to the invaded range samples (239.00 mm, 116.40 - 440.60 mm; 35.00 mm, 6.40 - 79.30 mm). Precipitation is additionally more variable in the native range, (native mean precipitation seasonality: 72.25, range: 36.64 - 116.40; invaded: 55.98, 31.22 - 103.68). The annual mean temperature of the invaded samples and native range samples were similar but there was higher variability in the native range (native mean annual temperature: 20.36 , range: 12.49 - 27.75 ; invaded: 19.27 , 14.63 - 22.95 ), while samples from the invaded range experienced more variable monthly temperatures than samples in the native range (invaded mean diurnal range: 12.34 , range: 9.68 - 15.62; native: 8.36 , range: 5.25 - 12.07 ).

Of the 4,291 variant loci retained in the dataset, there were 114 unique candidate SNPs associated with the five retained environmental variables tested (BIO1: annual mean temperature, BIO2: mean diurnal range, BIO13: precipitation of the wettest month, BIO14: precipitation of the driest month, and BIO15: precipitation seasonality) across the invaded range (Fig. 3). The greatest number of candidate SNPs were associated with precipitation of the driest month (BIO14; 36 SNPs), followed by mean diurnal range (BIO2; 27 SNPs), precipitation seasonality (BIO15; 18 SNPs), annual mean temperature (BIO1; 17 SNPs), and precipitation of the wettest month (BIO13; 16 SNPs). Among these 114 candidate SNPs there were 50 present in coding loci and 64 present in noncoding loci. There was no significant difference in the proportion of SNPs explained by each bioclimatic variable in coding versus noncoding loci (Chi-squared test; *P* = 0.9262, ^2^ = 0.88833, *df* = 4).

**Figure 3.**
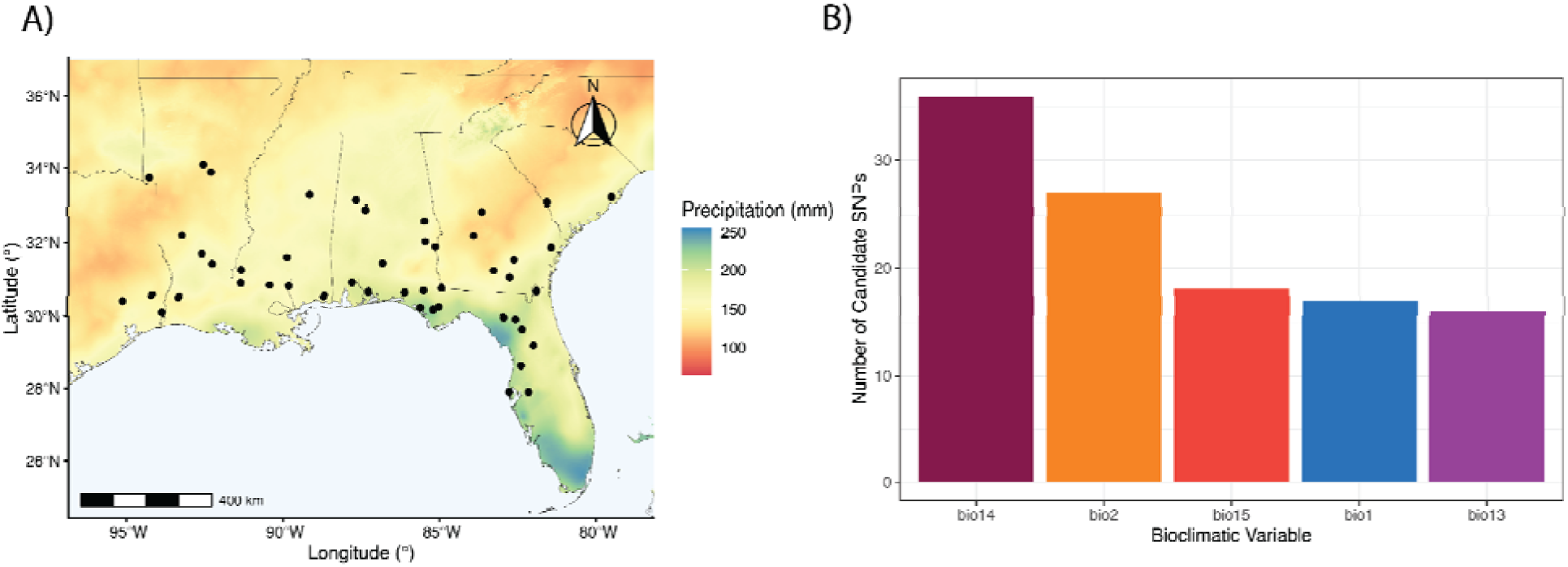
(A) An example environmental gradient (BIO14: precipitation of the driest month) in the invaded range of *L. japonicum*. Sample localities are depicted by black dots. (B) Descending bar graph depicting the number of candidate SNPs associated with five bioclimatic variables. BIO14: precipitation of driest month, BIO2: mean diurnal range, BIO13, precipitation of wettest month, BIO1: annual mean temperature, BIO15: precipitation seasonality.

### Changes in Genetic Variation Over Time

For non-coding loci, a model with an interaction term between invasion status and collection year was marginally significant (F_3,70_, *P* = 0.050, Adjusted R^2^ = 0.066), had a marginally significant interaction term (*P*=0.0511; Table S4) with a significant effect of invasion status (*P* = 0.0156) such that invaders had lower observed heterozygosity than native samples (β = -0.027; Fig. 4A). Given the significant interaction term, linear models were fit for each range independently. There was no significant association between collection year and observed heterozygosity in the native range (F_1,22_, *P* = 0.986, Adj. R^2^ = -0.045), but there was a significant, positive relationship in the invaded range (F_1,48_, β=2.978 × 10^−4^, *P* = 0.008, Adj. R^2^ = 0.1208; Table 1). At the beginning of the invasion (1940), the predicted observed heterozygosity was approximately 15% lower in the invaded range (0.152 vs 0.179 for the native range), whereas values were similar between ranges for the later samples from the invasion (Fig. 4A). For coding loci, the overall model with an interaction term was significant (F_3,70_, P = 0.049, Adj. R^2^ = 0.067), although no individual component was significant (all *P* > 0.05). Given that the interaction between invasion status and collection year was not significant (*P* = 0.266), a reduced linear model without an interaction term was fit, which was significant (F_2,71_, *P* = 0.036, Adj. R^2^ = 0.063) with a significant, positive relationship between collection year and observed heterozygosity (β = 2.483 × 10^−4^, *P* = 0.01) and no difference between native and invaded range samples (*P* = 0.327; Fig. 4B). For the GEA candidate loci (Fig. 4C), models were not significant fits either with (F_3,70_, *P* = 0.106, Adj. R^2^ = 0.044) or without (F_2,71_, *P*=0.999, Adj. R^2^ = -0.028) interaction terms.

**Figure 4.**
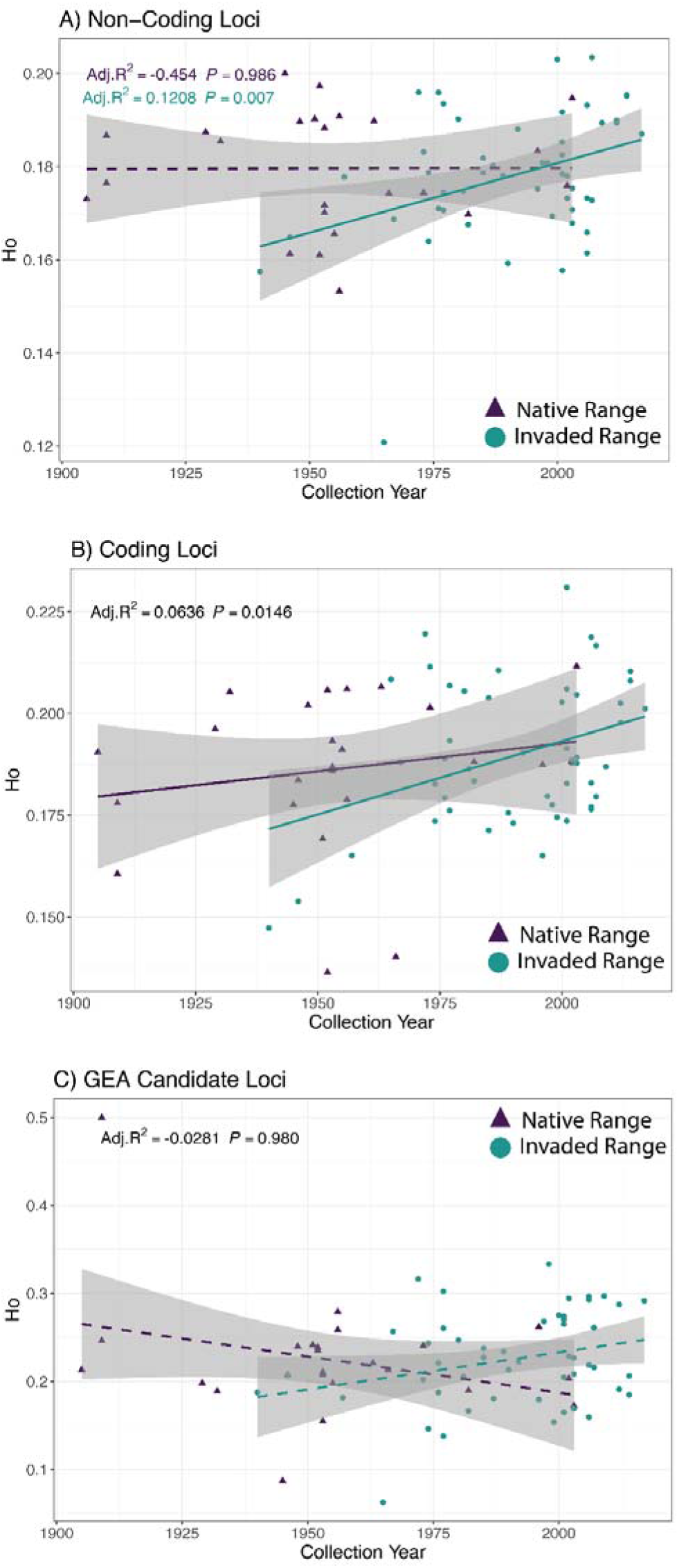
Temporal variation in individual heterozygosity in native (purple) and invaded (teal) ranges of *Lygodium japonicum* in (A) non-coding loci, (B) coding loci, and (C) GEA candidate loci. Note: y-axis scales differ between the three loci types. Solid lines indicate significant relationships and dashed lines indicate that relationships were non-significant.

**Table 1.** Summary of selected linear models testing the effect of collection year and invasion status on individual heterozygosity in coding loci, non-coding loci, and GEA candidate loci. Results for all linear models can be found in ‘Results’ and Table S4.

| **Model** | Non-coding Heterozygosity |  |  |  |  |  |
|---|---|---|---|---|---|---|
|  | **Effect** | **Estimate** | **SE** | ***t*** | ***P*** | **Adjusted *R*<sup>2</sup>** |
| Native | Collection Year | 1.821e-6 | 1.019e-4 | 0.018 | 0.9859 | -0.0454 |
| Invaded | Collection Year | 2.978e-4 | 1.071e-4 | 2.781 | **0.0073** | 0.1208 |
| Full Model (Both Ranges) Without Interaction Term | Coding Heterozygosity |  |  |  |  |  |
|  | Collection Year | 2.483e-4 | 9.917e-5 | 2.504 | **0.0146** | 0.0636 |
|  | Invasion Status | -5.627e-3 | 5.700e-3 | -0.987 | 0.3269 |  |
|  | GEA Candidate Heterozygosity |  |  |  |  |  |
|  | Collection Year | 8.53E-06 | 3.42E-04 | 0.025 | 0.98 | -0.0281 |
|  | Invasion Status | -2.56E-04 | 1.96E-02 | -0.013 | 0.99 |  |

*Note*: SE = 1 standard error

## DISCUSSION

We leveraged low-coverage whole genome resequencing data generated from a chronosequence of herbarium specimens collected between 1905-2017 to compare genomic variation and the capacity for rapid adaptation between the native and invaded range of the fern, *Lygodium japonicum*, in the southeastern U.S. Specifically, we analyzed population structure to attempt to identify source population(s) of the invasion, identified genetic variants that might be involved in adaptation to local climate conditions in the invaded range, and evaluated how individual-level genomic variation changed through time in the invasion. We found evidence of rapid adaptation to climate in the invaded range. We also found evidence of a genetic bottleneck in the invasion for non-coding variation but not for potentially adaptive variation, highlighting the maintenance of the capacity for rapid adaptation during invasion.

To date, there have been no investigations into the phenotypic basis of local adaptation to climate in the native or invaded ranges of *Lygodium japonicum*. Previous work investigating local adaptation in fern sporophytes and gametophytes suggests that growth rate and plant size are implicated in local adaptation (Hsu et al., 2014), as well as gametophyte senescence timing (Chambers and Emery, 2016). Understanding what traits are adaptive in different environmental conditions in *L. japonicum* could provide insight into a variety of relevant ecological and evolutionary processes, including which traits may have facilitated the *L. japonicum* invasion, and which traits may be exploited for managing invasive populations.

### The Origin(s) of the Lygodium japonicum Invasion Remain Unclear

In a previous analysis of chloroplast genomes from this dataset, Markley et al. (2024) found that one rare haplotype from the invasion was shared with samples from China and the Philippines, but the dominant haplotype from the invasion was not observed in the native range samples. Although the dominant invader haplotype was most similar to one from China, a definitive assignment of the invaders to one or more regions in the native range was not possible. Using genome-wide nuclear data, we recovered similar patterns but were ultimately unable to further clarify the origin of the *L. japonicum* invasion. Four individuals from the native range clustered with samples from the invaded range, all of which were from China or Indonesia. Notably, these four native samples clustered with samples from early in the invasion, suggesting that the genetic makeup of the earliest specimens from the invasion were indeed most similar to the native range. However, other native individuals from China and Indonesia did not cluster with the invaded range samples, making it difficult to pinpoint either of these as putative source regions. We also found that the samples with rare chloroplast haplotypes in the invasion were not significantly more distant in nuclear content than samples with the dominant haplotype were from one another, suggesting that these chloroplast lineages had admixed in the invasion by the time of sampling. Relatedness among individuals determined from nuclear and plastid genomic data will often conflict when there is a history of recent admixture or chloroplast capture (Rieseberg and Soltis, 1991).

The introduction history of the *L. japonicum* invasion could explain the admixture of divergent lineages and the lack of a close relationship with native genotypes. *Lygodium japonicum* was introduced to the U.S. through the horticultural trade (Koop, 2009), from which it escaped and became naturalized, a common route of colonization for ferns (van Kleunen et al., 2018; McCulloch-Jones et al., 2021). Horticultural practices can involve breeding individuals from multiple source regions in nurseries. Tracing the history of source material that was used to create these artificial crosses can be difficult, especially with older nurseries that may not have kept records of their source material (e.g., Arias and Rieseberg, 1995). Such breeding processes could potentially blur signals of ancestry in commercial stock that eventually escaped into natural ecosystems (Culley and Hardiman, 2009; Holt and Beck, 2023). Using population-level genomic sampling, models of introduction sources and admixture could be tested (Dlugosch et al., 2015; Welles and Dlugosch, 2019), which would reveal insights into how horticultural selections were made and potential combined in the history of *L. japonicum* cultivation.

### Rapid Adaptation to Climate in the Invasion

Genotype environment analysis identified114 unique, putatively adaptive genomic loci associated with the five tested bioclimatic variables (Figure 3B). Most candidate loci were significantly associated with precipitation of the driest month (BIO14; Figure 3B), indicating that water limitation may be an important source of selection on invasive *L. japonicum* populations. This is consistent with evidence from biogeographical investigations that find that fern distributions are shaped by water availability as well as temperature (Khine et al., 2019; Suissa et al., 2021; Shan et al., 2025). Water availability has long been thought to act as a strong selective pressure on ferns in part because they have motile sperm that require water to swim to eggs for fertilization (Haufler et al., 2016), Indeed, Grusz et al. (2021) posit that drought and other abiotic conditions have driven the repeated evolution of apomixis in xeric-adapted ferns, to avoid the need for fertilization via swimming sperm. Our results suggest that introduced *L. japonicum* populations have been successfully adapting to this key axis of environmental variation in their invasion, though changes at dozens of loci across the genome.

Interestingly, mean diurnal range (BIO2) was associated with many candidate candidate loci, suggesting that the degree of daily temperature fluctuations may impose selection on *L. japonicum* populations as well. Models of the *L. japonicum* distribution in its native range indicate that mean diurnal range is an important driver of its distribution (Shan et al., 2025). The magnitude of daily and seasonal variation in temperature can influence the ability of individuals to reach maximum photosynthetic rates (Farquhar et al., 1980; Volkova et al., 2011; Krieg and Chambers, 2022), affecting individual growth and performance when fluctuations exceed the amount of variation to which an individual can effectively respond. This is particularly apparent if an individual experiences temperature beyond its physiological limit during a large temperature fluctuation (e.g., freezing; Lott et al., 2003; Pelosi et al., 2025). Again, our results indicate that *L. japonicum* populations have been successful in adapting to variation in this important aspect of their environment through variants at multiple loci.

Putative functions were assigned to twelve of the candidate loci by identifying *Arabidopsis thaliana* genes with significant sequence similarity (e.g., putative orthologs). Specifically, six unique, similar *A. thaliana* mitochondrial and nuclear genes were identified that are involved in a variety of functions (Table S5). Notably, the pentatricopeptide repeat (PPR) protein coded by the *DOT4* gene (AT4G18750.1) has high sequence similarity to a locus with a SNP in our dataset associated with precipitation of the wettest month (BIO13) and is known to be involved in *A. thaliana* leaf and root development (Hayes et al., 2013). In ferns, the PPR family is associated with extensive RNA editing (Wolf et al., 2004), which may be adaptive (Fauskee et al., 2025). If and how proteins in this family are associated with climate adaptation in ferns necessitates further investigation. Additionally, the cysteine-rich receptor-like protein kinase coded by the *CRK8* gene (AT4G23160.2) has significant sequence similarity to four loci with GEA candidate SNPs that are associated with precipitation of the wettest month (BIO13) and precipitation seasonality (BIO15). In *A. thaliana*, *CRK8* has an upstream retrotransposon insertion that modulates a defense-stress tolerance tradeoff (Macquet et al., 2025). Tradeoffs between defense and tolerance functions are hypothesized to be an important factor in defining successful strategies of invasive species (Mayrose et al., 2011; Mesa and Dlugosch, 2020). *CRK* genes code for cell membrane proteins that are involved in a range of both immune and abiotic stress responses (Zhang et al., 2023), and characterizing their role in ferns could possibly provide insight into the mechanisms underlying drought responses, abiotic stress tolerance, and resource allocation to defense and stress in both invasive and native ferns.

### Evidence for Loss of Non-Coding, but not Adaptive, Genetic Variation

Researchers have long pondered how invasive populations undergoing massive reductions in numerical size during introduction could overcome associated genetic bottlenecks (Baker and Stebbins, 1965). However, Nei et al. (1975) found that if population growth is fast and several individuals (e.g. as few as ten under realistic conditions) make up the founding population, the reduction in heterozygosity is low even in the face of population bottlenecks. Both of these criteria are common in invasive plants, which tend to have rapid growth and are not typically introduced as individuals (e.g., as in the horticultural trade where rare plants are imported in batches of dozens or more (Naqinezhad et al., 2024). Furthermore, if population growth is fast, loss of heterozygosity is lost only in the early generations. The question of how much variation is established and how it affects the ability of introduced populations to adapt has nevertheless persisted into the genomic era (Hodgins et al., 2025).

Using genomic data from samples taken across the invasion, decreased heterozygosity in non-coding loci in the invaded range was observed in the earliest samples. Although these samples have ∼15% lower heterozygosity than native range individuals, these values increased significantly over time and reached the same values as natives by the later samples (after the year 2000) in the invaded range, indicating a genetic bottleneck that is small in magnitude and typical of introduced species as a whole (Dlugosch and Parker, 2008). However, only the earliest samples have individual heterozygosity lower than in the native range, with several samples reaching individual levels of heterozygosity above the native range as early as the 1970s. This rapid increase in heterozygosity suggests multiple introductions may have restored genetic variation, consistent with our inferences of introduction history. Notably, the time ranges for which we have samples from each range is different (1905-2003 for native; 1940-2007 for invasive), which might overstate direct comparisons in the full models. However, subsampling to include only samples with overlapping time ranges would limit the power of such an analysis.

The *type*, rather than simply the amount, of genetic variation retained during colonization is critical to understanding the capacity of invasive species for rapid adaptation (Dlugosch et al., 2015). Indeed, overall and adaptive variation are not always correlated, because selection will shape the retention or loss of adaptive variation (e.g., Marila and Crnokrak 2001; McKay and Latta, 2002; Leionen et al., 2008). In the *L. japonicum* invasion we found that, while non-coding loci had a significant decrease in heterozygosity in the invaded range, there was no difference in the trends of individual heterozygosity in coding loci, which are more likely to be under selection and contain adaptive variants (e.g., Williamson et al., 2014). Furthermore, putatively adaptive loci associated with climatic variables (GEA candidates) showed no difference in heterozygosity between the two ranges and there was not a significant increase in heterozygosity over time in either range. Thus no evidence was seen for potentially adaptive variation experiencing the same genetic bottleneck as was observed in non-coding (putatively neutral) variation or as Markley et al. (2024) observed in chloroplast haplotype variation. This suggests that selection may have favored the maintenance of potentially adaptive variants during invasion. Importantly, invaded range *L. japonicum* individuals are autopolyploid at least in part of the range (Pelosi et al., 2023) , which may facilitate the preservation of genetic diversity (te Beest et al., 2012).

## Conclusion

We found that trends of genetic diversity over the course of an invasion differ depending on the type of variation being analyzed. In particular, the demographic bottleneck in the *L. japonicum* introduction appears to be associated with a small genetic bottleneck in putative neutral variation, but no bottleneck in potentially adaptive variation. Our findings add support to the idea that invaders may often establish ample genetic variation for rapid adaptation, including the under-studied invasive ferns.

## DATA AVAILABILITY

All genomic sequencing data used for our analyses are publicly available on NCBI; accession numbers for each individual used in the present study are listed in Table S1. A GitHub repository (https://github.com/apearse22/LYJA_invasion_genomics) contains all of the code used in our analyses.

## ACKNOWLEDGEMENTS

We thank Katie Eaton for her advice on statistical methodology. This work was supported by USDA NIFA Postdoctoral Research Fellowship #2024-67012-43394 to J.A.P. Participation by A.R.P. was supported by the University of Arizona Undergraduate Biology Research Program with funding from the University of Arizona Office of Research & Partnerships and USDA NIFA #2024-67012-43394. K.M.D. and P.C.N. were supported by USDA NIFA #2023-67013-40169. Sequence data were generated with funding from NSF OIA #1920858 to J.B.B.

## SUPPLEMENTAL MATERIAL

**Supplemental Table 1.** Associated metadata with the samples used in this study.

**Supplemental Table 2.** Mantel test results for isolation by geographic and temporal distance.

**Supplemental Table 3.** Individudal-level observed heterozygosity (Ho) for each sample in our dataset.

**Supplemental Table 4**. Results of linear models for non-coding, coding, and GEA candidate loci.

**Supplemental Table 5.** GEA candidate loci with BLAST hits to *Arabidopsis thaliana* (A. th) genes and their functions.

**Supplemental Figure 1.** Environmental gradients of the five selected bioclim variables (BIO1, BIO2, BIO13, BIO14, and BIO15). Points are sample collection locations.

**Supplemental Figure 2.** Histogram (black) showing the median genetic distances for each individual in the major chloroplast haplotype (haplotype #1). Red vertical line is the median genetic distance for the one sample with plastid haplotype #2 and plastid haplotype #1 individuals. Blue vertical lines are median genetic distances for the two samples with plastid haplotype #3 and plastid haplotype #1 individuals.

## Notes

### Competing Interest Statement

The authors have declared no competing interest.

